## Supplemental Data for "Inhibition of the Notch signal transducer CSL by Pkc53E-mediated phosphorylation to fend off parasitic immune challenge in *Drosophila*"

#### Supplemental Materials

#### Figures Supplement

**Figure 1 - Figure supplement 1**

*Notch acts upstream of Su(H); minor changes in hemocyte numbers in Su(H)$^{S269}$ phospho-mutants*

**Figure 1 - Figure supplement 2**

*Representative images for the various settings*

**Figure 2 - Figure supplement 1**

*The α-pS269 antiserum detects the phospho-mimetic Su(H) variant in vitro*

**Figure 3 – Figure supplement 1**

*Overview of the results from the kinase screen*

**Figure 3 – Figure supplement 2**

*MS/MS spectra of the phosphorylated Su(H) peptide*

**Figure 4 – Figure supplement 1**

*Conservation of Pkc53E and generation of an activated Pkc53E$^{EDDD}$ isoform*

**Figure 5 – Figure supplement 1**

*Expression of Su(H)$^{VP16}$-myc is not influenced by PMA or STAU*

**Figure 6 – Figure supplement 1**

*Representative images for the various settings.*

**Figure 6 – Figure supplement 2**

*The Pkc53E$^{\Delta 28}$ allele is a null mutant*

**Figure 7 – Figure supplement 1**

*Representative images for the various settings.*

**Figure 7 – Figure supplement 2**

*Raw blots of co-IP*

**Figure 8 – Figure supplement 1**

*Pkc53E is expressed in hemocytes.*

**Figure 9 – Figure supplement 1**

*Pkc53E$^{\Delta 28}$ is sensitive to wasp infestation*

#### Supplemental Materials

##### Figures Supplement

**Figure 1 - Figure supplement 1**

*Notch acts upstream of Su(H); minor changes in hemocyte numbers in Su(H)$^{S269}$ phospho-mutants*

**Figure 1 - Figure supplement 2**

*Representative images for the various settings*

**Figure 2 - Figure supplement 1**

*The α-pS269 antiserum detects the phospho-mimetic Su(H) variant in vitro*

**Figure 3 – Figure supplement 1**

*Overview of the results from the kinase screen*

**Figure 3 – Figure supplement 2**

*MS/MS spectra of the phosphorylated Su(H) peptide*

**Figure 4 – Figure supplement 1**

*Conservation of Pkc53E and generation of an activated Pkc53E$^{EDDD}$ isoform*

**Figure 5 – Figure supplement 1**

*Expression of Su(H)$^{VP16}$-myc is not influenced by PMA or STAU*

**Figure 6 – Figure supplement 1**

*Representative images for the various settings.*

**Figure 6 – Figure supplement 2**

*The Pkc53E$^{\Delta 28}$ allele is a null mutant*

**Figure 7 – Figure supplement 1**

*Representative images for the various settings.*

**Figure 7 – Figure supplement 2**

*Raw blots of co-IP*

**Figure 8 – Figure supplement 1**

*Pkc53E is expressed in hemocytes.*

**Figure 9 – Figure supplement 1**

*Pkc53E$^{\Delta 28}$ is sensitive to wasp infestation*

**Figure 9 – Figure supplement 2**

*Representative images for the various settings.*

#### Tables Supplement

**Figure 3 –Table 1 supplement**

List of kinases predicted to recognize S269 in Su(H) as substrate *in silico*

**Figure 3 –Table 2 supplement**

List of kinases accepting the BTD domain of Su(H) as substrate *in vitro*

**Figure 3 –Table 3 supplement**

Fly strains used for the larval crystal cell screen

**Figure 3 –Table 4 supplement**

Larval crystal cell screen

#### Supplemental Tables, Materials and Methods

**S1 Table** Key resources

**S2 Table** Oligonucleotides

**Source data**

**Figure 1 source data**

*Raw data and statistical analysis*

**Figure 2 source data**

*Raw data and statistical analysis*

**Figure 4 source data**

*Raw data and statistical analysis*

**Figure 6 source data**

*Raw data and statistical analysis*

**Figure 7 source data**

*Raw data and statistical analysis*

**Figure 9 source data**

*Raw data and statistical analysis*

### Figures Supplement

#### *Figure 1 - Figure supplement 1*

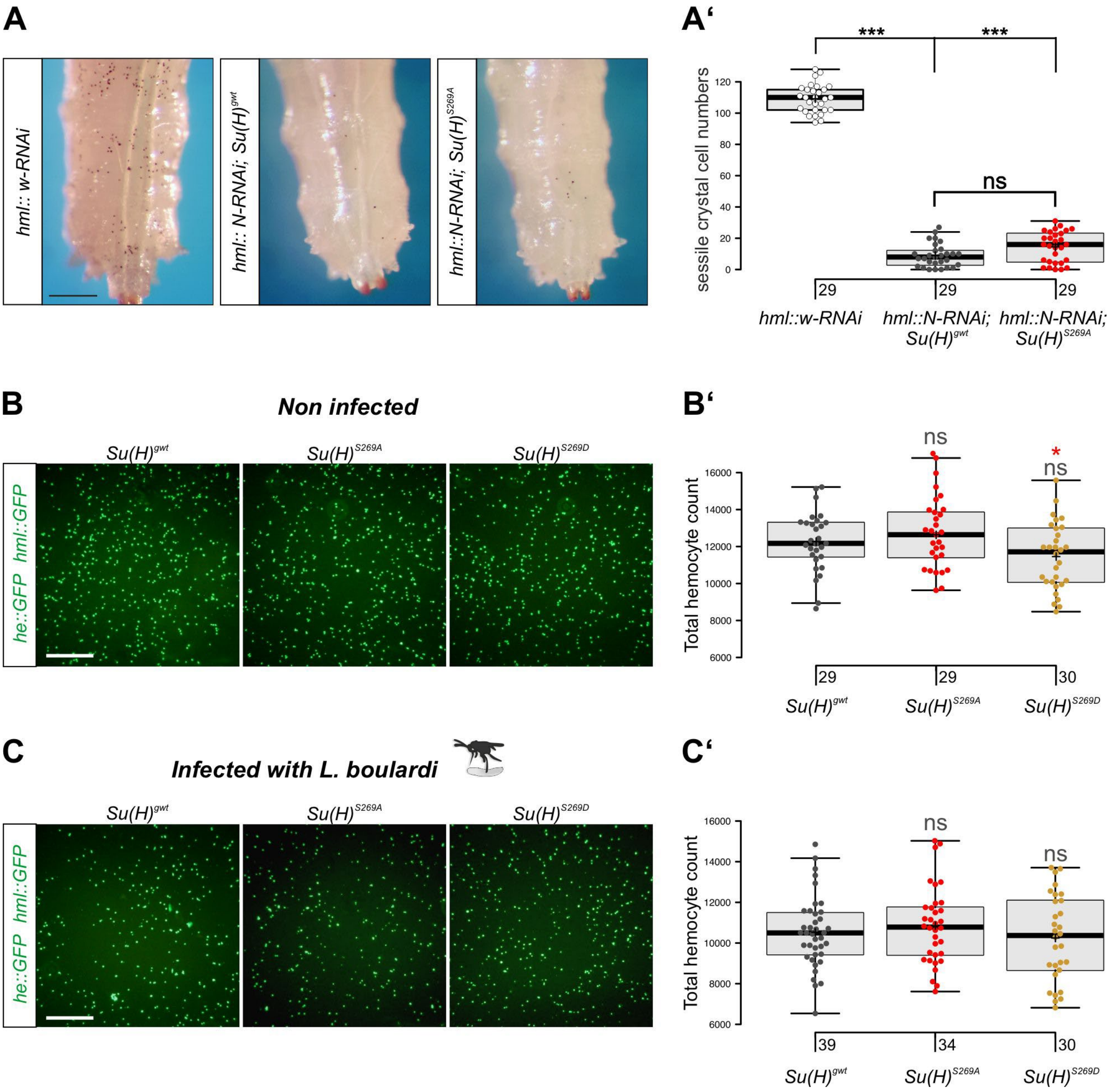

***Notch acts upstream of Su(H); minor changes in hemocyte numbers in $Su(H)^{S269}$ phospho-mutants***
**(A,A')** *Notch-*$Su(H)^{S269A}$ epistasis experiment. RNAi against *Notch* was induced in the hemocytes with *hml*-Gal4 (*hml*::*N*-RNAi) in either $Su(H)^{gwt}$ or $Su(H)^{S269A}$ background; *hml*::*white*-RNAi served as control. (A) Representative pictures of larvae are shown. Scale bar, 250 µm. (A') Crystal cell numbers were determined in the last two segments of heated third instar larvae (each dot represents one larva; n, as shown)**.** Note the strong drop in crystal cell numbers in the *hml*::*N*-RNAi larvae compared to control *hml*::*white*-RNAi. There is no significant difference between the $Su(H)^{gwt}$ or $Su(H)^{S269A}$ background. Statistical analyses with Kruskal-Wallis test, followed by Dunn's multiple comparison test with *** $p<0.001$ and ns (not significant $p\geq0.05$). **(B-C')** Total hemocyte count in uninfected

larvae (B,B') or in larvae 24 h after infection with *L. boulardi* (C,C'). (B,C) Hemocytes were visualized by green fluorescence (*he*::GFP *hml*::GFP). Representative pictures of hemolymph are shown. Scale bar 300 µm. (B',C') The total number of hemocytes in $Su(H)^{gwt}$, $Su(H)^{S269A}$ or $Su(H)^{S269D}$ larvae in each condition was not significantly different, albeit a subtle increase in $Su(H)^{S269A}$ and a subtle decrease in $Su(H)^{S269D}$ was noted. Each dot represents the hemocyte count of one larva; the total number of animals tested is indicated below. Statistical analysis with ANOVA, followed by Tukey's multiple comparisons test relative to control $Su(H)^{gwt}$, * $p<0.05$, ns (not significant $p\geq0.05$).

*Figure 1 - Figure supplement 2*

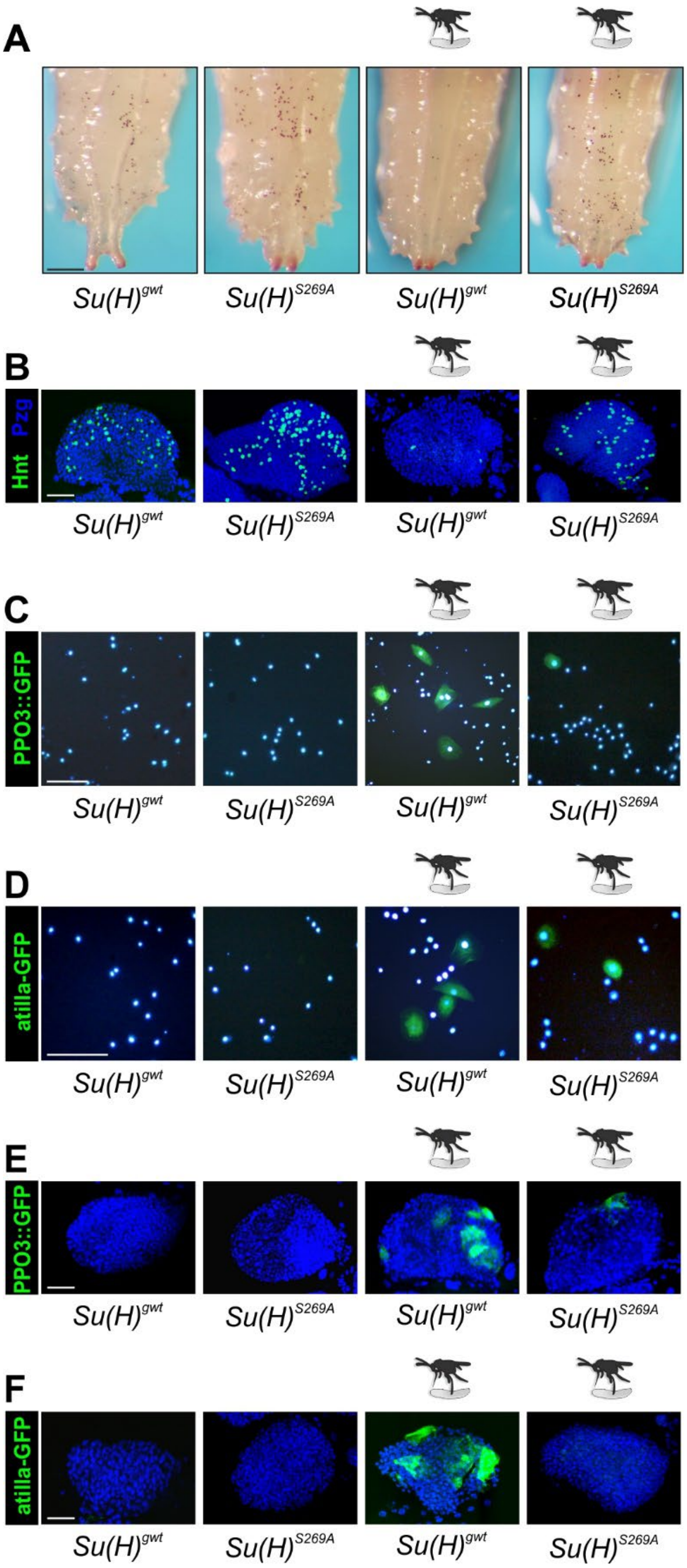

***Representative images for the various settings***

**(A)** Representative images of heated larvae of the given genotype used for crystal cell counting.

**(B,E,F)** Representative images of one lymph gland 1° lobe. Nuclei stained with Pzg antibodies (blue). Labelling of crystal cells (Hnt in B), or of lamellocytes (*PPO3*::GFP in E, *atilla*-GFP in F) is shown in green. **(C,D)** Representative images of hemolymph derived from larvae of the given genotype. Lamellocytes labelled green with *PPO3*::GFP (C) or *atilla*::GFP (D); nuclei blue (DAPI). (A-F) Infestation with *L. boulardi* indicated by the wasp schematic. Scale bar: 250 µm in (A), 50 µm in (B-F).

***Figure 2 – Figure supplement 1***

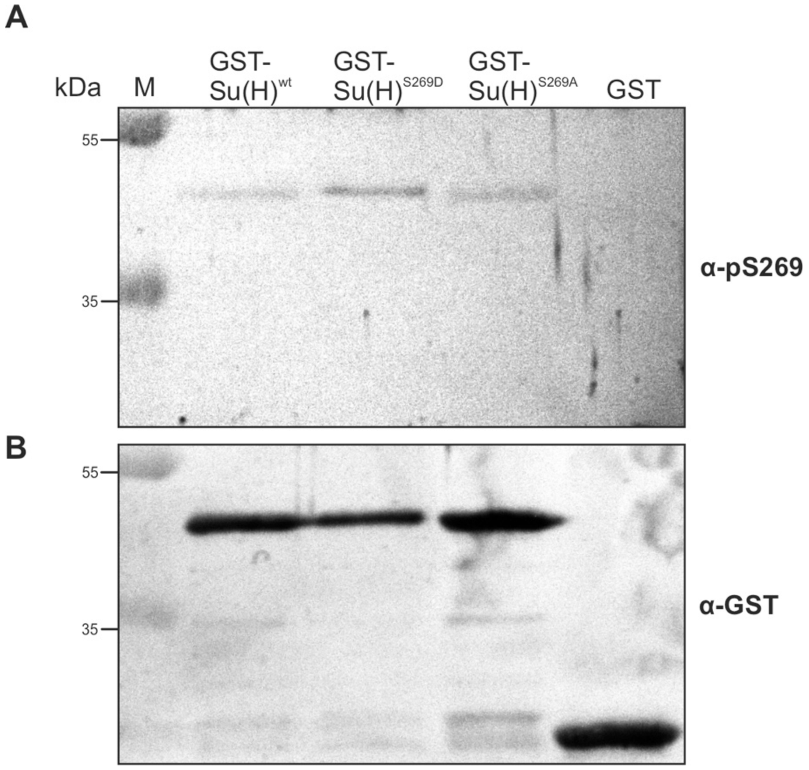

***The α-pS269 antiserum detects the phospho-mimetic Su(H) variant in vitro***

(**A**) Western Blot with purified GST-BTD-Su(H) phospho-variants as indicated, as well as GST alone as control. pS269 antibody detects preferentially the phospho-mimetic BTD-Su(H)$^{S269D}$ variant, and to a lesser degree wild type BTD-Su(H)$^{wt}$ as well as BTD-Su(H)$^{S269A}$. (**B**) Same samples detected with α-GST. Note unequal loading: there is more GST-Su(H)$^{wt}$ and GST-Su(H)$^{S269A}$ proteins loaded in comparison to BTD-Su(H)$^{S269D}$.

*Figure 3 – Figure supplement 1*

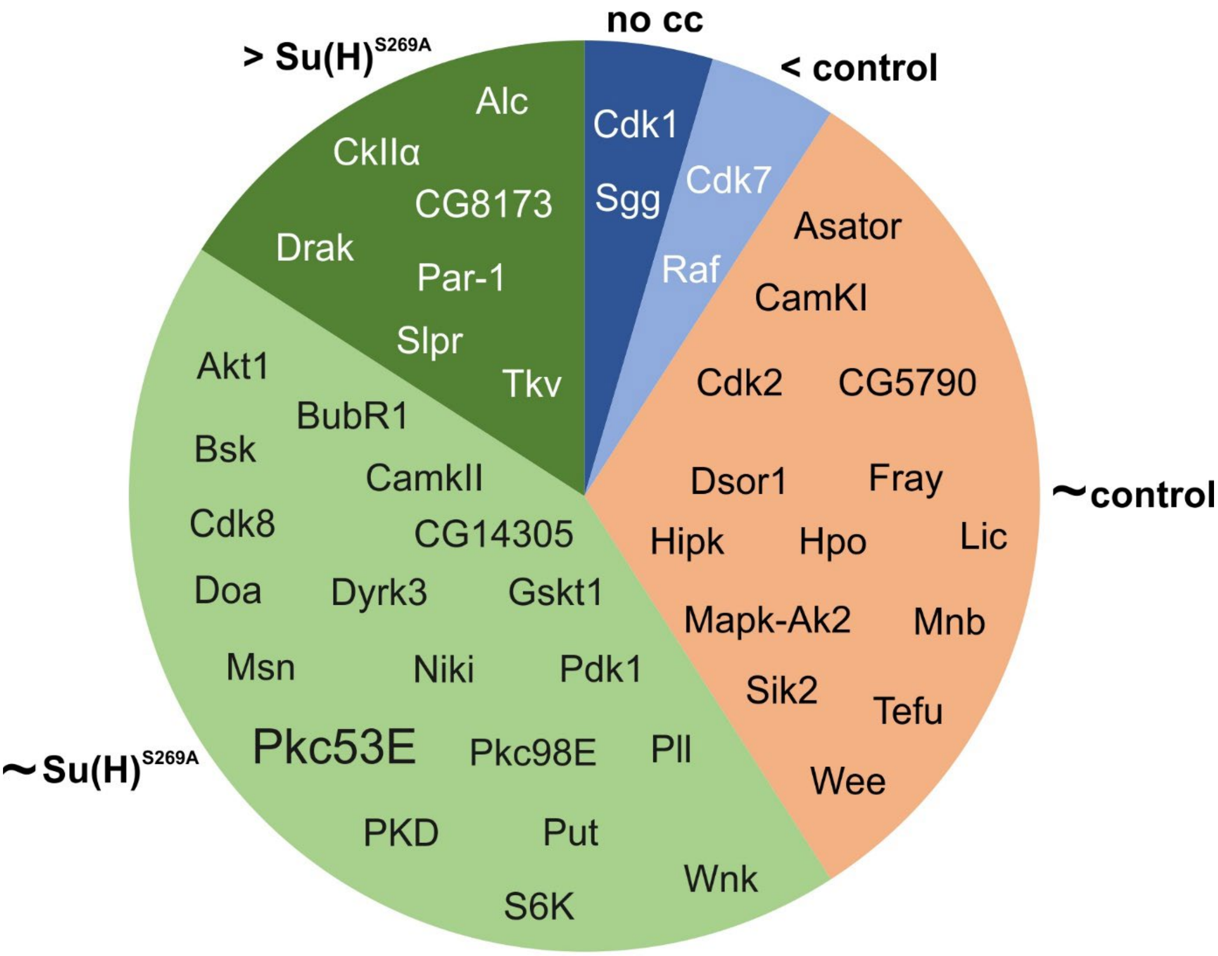

***Overview of the results from the kinase screen***

Crystal cell numbers recorded in the respective kinase mutants served the classification. Dark blue: no crystal cells; light blue: reduced number of crystal cells; orange: numbers match the control; light green: numbers match *Su(H)*$^{S269A}$ mutants; dark green: numbers largely exceed the *Su(H)*$^{S269A}$ mutants. More details are found in supplemental Table 4.

*Figure 3 – Figure supplement 2*

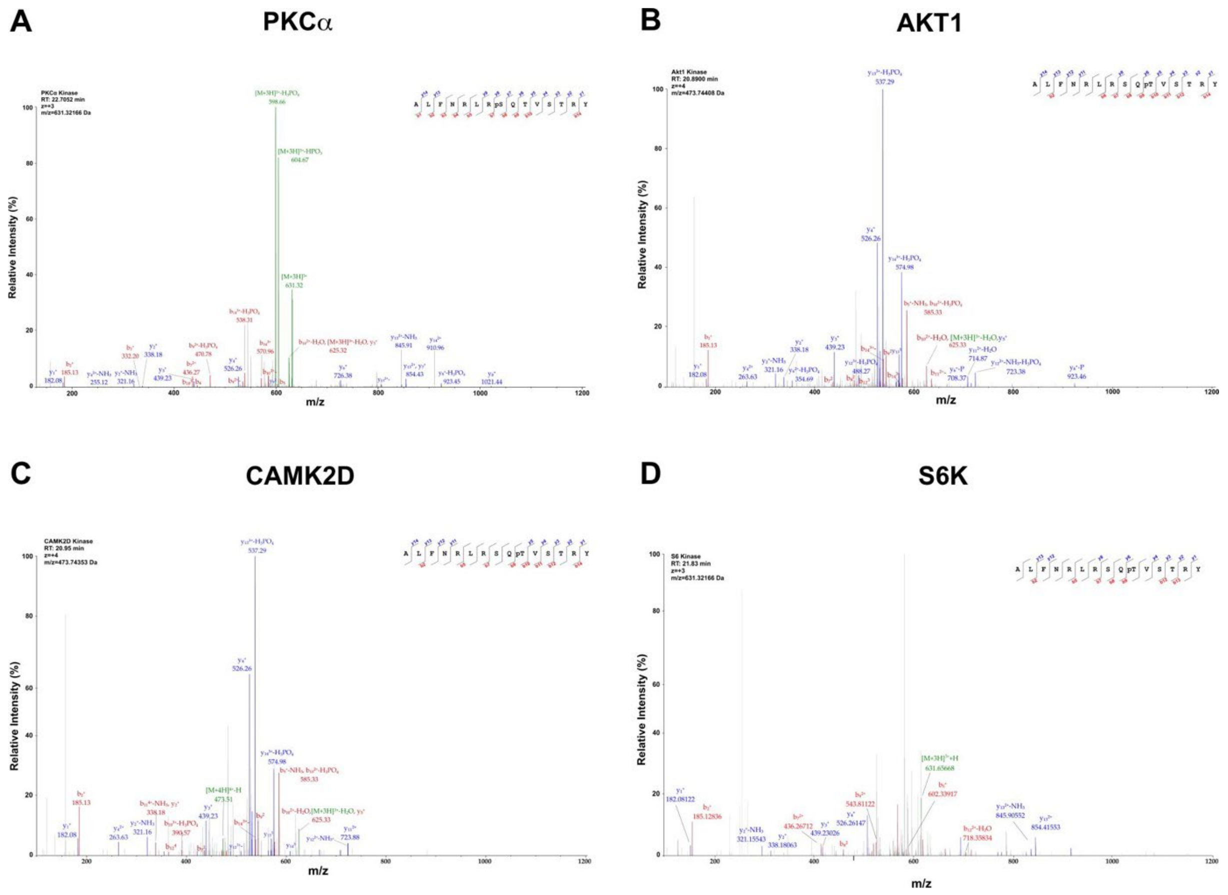

***MS/MS spectra of the phosphorylated Su(H) peptide***

**(A)** MS/MS spectrum on the Su(H) phosphopeptide 262-ALFNRLRpSQTVSTRY-276 after treatment with activated human kinases PKCα. (**B-D**) MS/MS spectra of the Su(H) phosphopeptide 262-ALFNRLRSQpTVSTRY-276 after incubation with activated human AKT1 (B), CAMK2D (C) and S6 Kinase (D), respectively. The phosphorylation at S8, corresponding to S269 and at T10, corresponding to T271 in Su(H) was confirmed by b-and y-ion series as indicated in blue and red, respectively. Neutral loss reactions of $H_2O$ and $H_3PO_4$ from the precursor peptide are indicated in green.

*Figure 4 – Figure supplement 1*

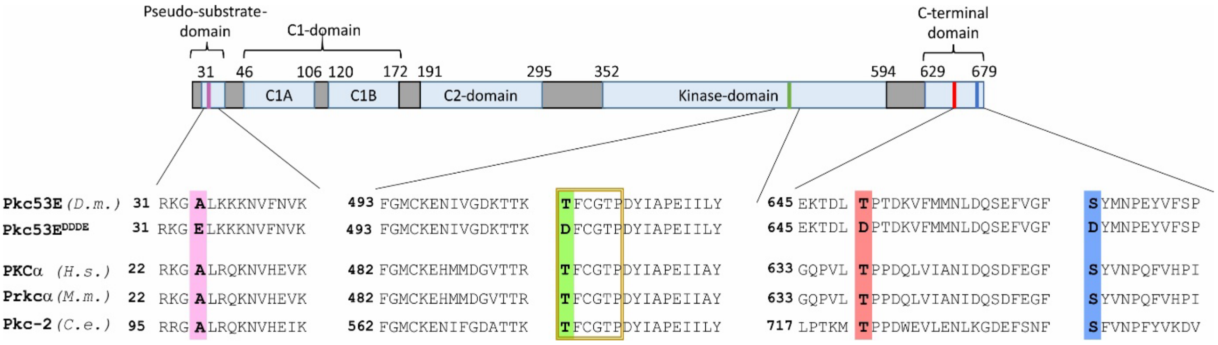

***Conservation of Pkc53E and generation of an activated Pkc53E$^{EDDD}$ isoform***

The typical PKC structure is shown, including the highly conserved pseudo-substrate domain at the N-terminus, the co-factor sensitive C1- and C2-domains, the catalytically active kinase domain and the C-terminal domain, harbouring the activation center. A comparison of these conserved domains including Pkc53E from *Drosophila melanogaster* (D.m.), PKCα from *Homo sapiens* (H.s.), Prkcα from *Mus musculus* (M.m.) and Pkc-2 from *Caenorhabditis elegans* (C.e.) is depicted. Highlighted in colour are the amino acids phosphorylated in the course of PKC activation, as well as their exchange to an aspartate (D) or glutamate (E) residue in the activated Pkc53E$^{EDDD}$ mutant isoform.

*Figure 5 – Figure supplement 1*

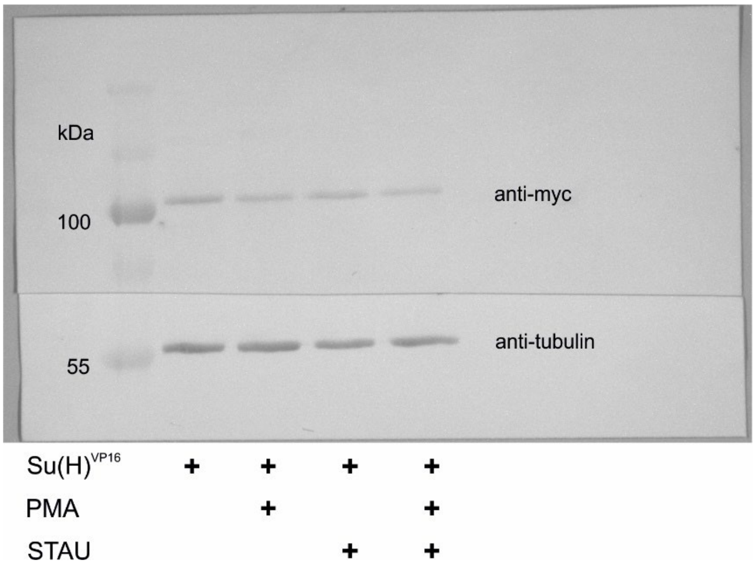

***Expression of Su(H)$^{VP16}$-myc is not influenced by PMA or STAU***

Western blot of HeLa RBPj$^{ko}$ cells, transfected with Su(H)$^{VP16}$-myc and treated with PMA and/or STAU as indicated. Expression of Su(H)$^{VP16}$ was detected with anti-myc antibodies; beta-tubulin expression served as loading control. The blot was sliced before independent treatment with the two antibodies; the entire blot is shown.

***Figure 6 – Figure supplement 1***

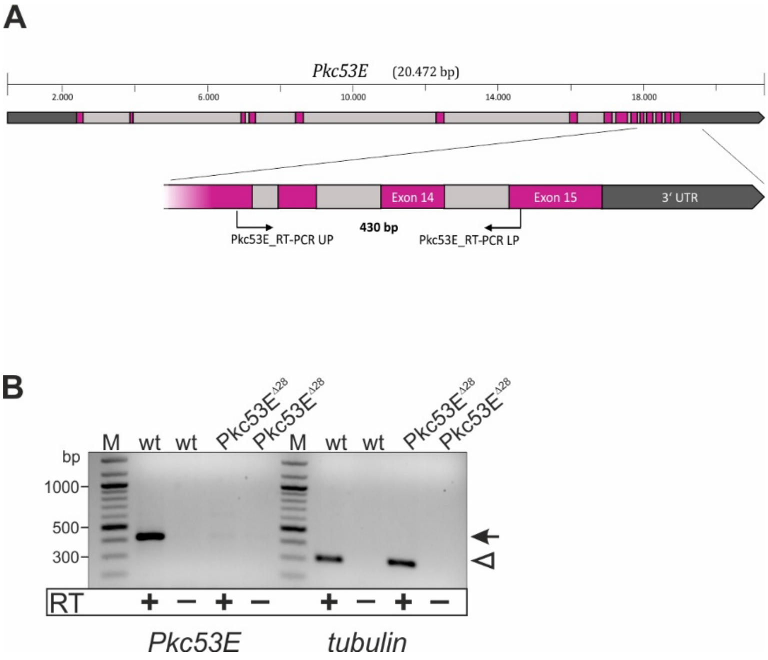

***The Pkc53E$^{\Delta28}$ allele is a null mutant***

(**A**) Genomic map of the *Pkc53E* locus covering in total more than 20 kilo bases (kb). Numbers above give the size of the whole locus on the right arm of the second chromosome. Exons are shown as magenta boxes, introns as light grey boxes; untranslated regions (UTR) in dark grey. The region amplified in the RT-PCR is enlarged underneath. Primers were chosen to overlap the last three introns of *Pkc53E.* (**B**) RT-PCR performed on RNA of each 50 *Pkc53E$^{\Delta28}$* and *$y^1$ $w^{67c23}$* control larvae (wt), respectively, with (+) and without (-) reverse transcriptase. Primer pairs *Pkc53E_RT-PCR UP/LP* overlap the last three introns of *Pkc53E*, resulting in a product of 430 bp (arrow). Tubulin served as control for intact mRNA (*tubulin*, expected product 299 bp, open arrowhead). M, size standard (1000 bp, 500 bp and 300 bp are labelled for reference). *Pkc53E* transcript was detected in the wt control, but not in *Pkc53E$^{\Delta28}$*, demonstrating the mutant to be null.

*Figure 6 – Figure supplement 2*

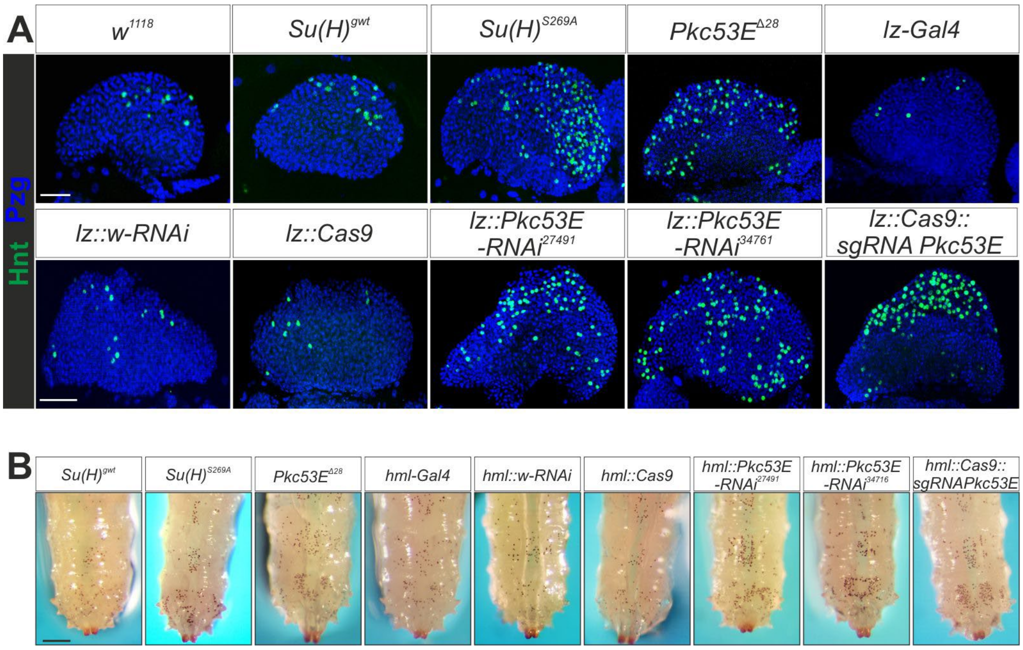

***Representative images for the various settings.***

**(A)** Representative images of one 1° lobe of the lymph gland derived from larvae of the given genotype. Crystal cells were stained with anti-Hnt (green), and nuclei with anti-Pzg (blue). Scale bar, 50µm. **(B)** Representative images of heat induced larvae of the given genotype used for crystal cell counts. Scale bar, 250 µm.

*Figure 7 – Figure supplement 1*

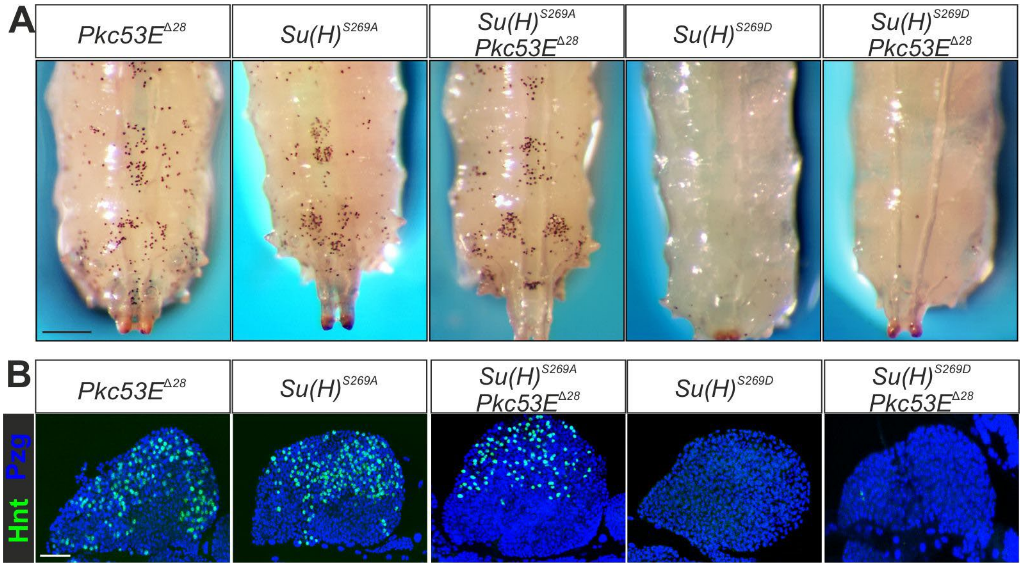

***Representative images for the various settings.***

**(A)** Representative images of heat induced larvae of the given genotype used for crystal cell counts. Scale bar, 250 µm. **(B)** Representative images of one 1° lobe of the lymph gland derived from larvae of the given genotype. Crystal cells were stained with anti-Hnt (green), and nuclei with anti-Pzg (blue). Scale bar, 50 µm.

*Figure 7 – Figure supplement 2*

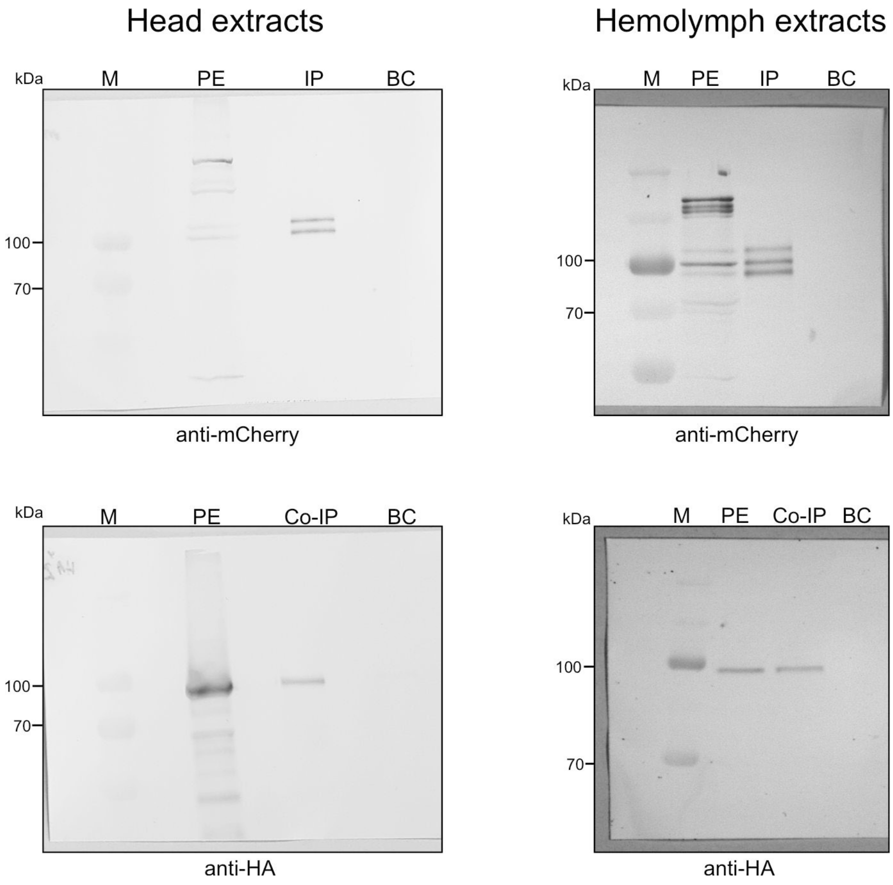

***Raw blots of co-IP***

RFP-Trap IP on protein extracts from heads (left blots) and hemolymph (right blots) derived from *gmr*::Pkc53E-HA Su(H)$^{gwt\text{-}mCh}$ or *hml*::Pkc53E-HA Su(H)$^{gwt\text{-}mCh}$ animals, respectively. Trapped Su(H)$^{gwt\text{-}mCh}$ was detected with anti-mCherry; co-precipitated Pkc53E-HA with anti-HA antibodies as indicated. PE, protein extract; IP, immune precipitate trapped; BC, binding control. M, pre-stained protein ladder; protein size is given in kDa.

*Figure 8 – Figure supplement 1*

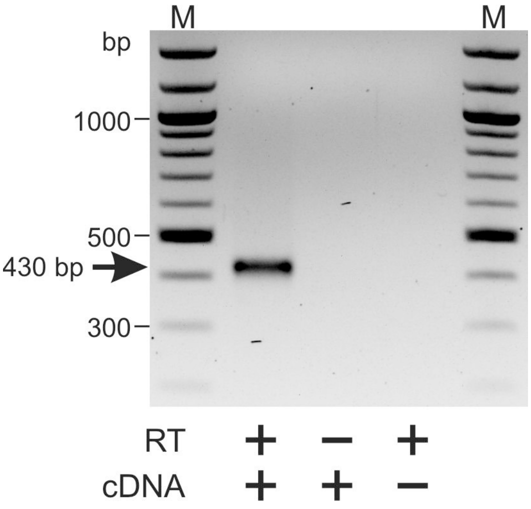

***Pkc53E is expressed in hemocytes.***

RT-PCR for *Pkc53E* expression in hemocytes of *Su(H)*$^{gwt}$ control larvae. A PCR-product of 430 bp is expected (arrow). RT, reverse transcriptase. M, 100 bp DNA-ladder.

*Figure 9 – Figure supplement 1*

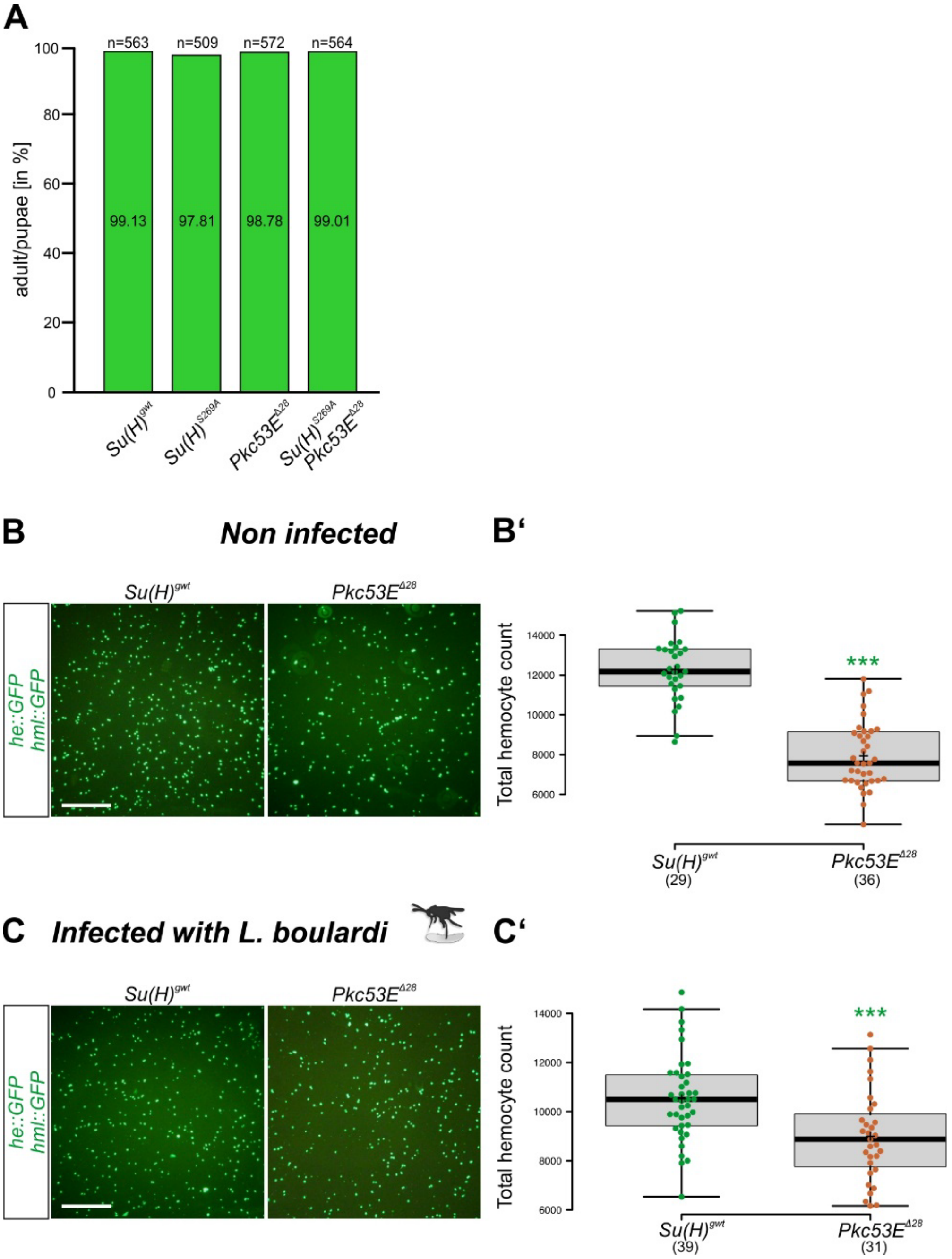

***Pkc53E$^{\Delta 28}$* is sensitive to wasp infestation.**

**(A)** Fly eclosure in the absence of wasp infestation. Percentage of adult flies with the given genotype hatched from at least 500 pupae (number as indicated). Statistical analyses with Kruskal-Wallis test, followed by Dunn's multiple comparison test, ns (not significant p≥0.05). **(B,C)** Representative pictures of hemocytes (green) labelled with *he*::GFP *hml*::GFP, not/infested with *L. boulardi* as indicated. Scale bar, 300 µm. (B',C') Every dot represents the hemocyte count of one larvae; number of analyzed larvae is shown below**.** Statistical analyses with unpaired Student's t-test, *** p<0.001.

*Figure 9 – Figure supplement 2*

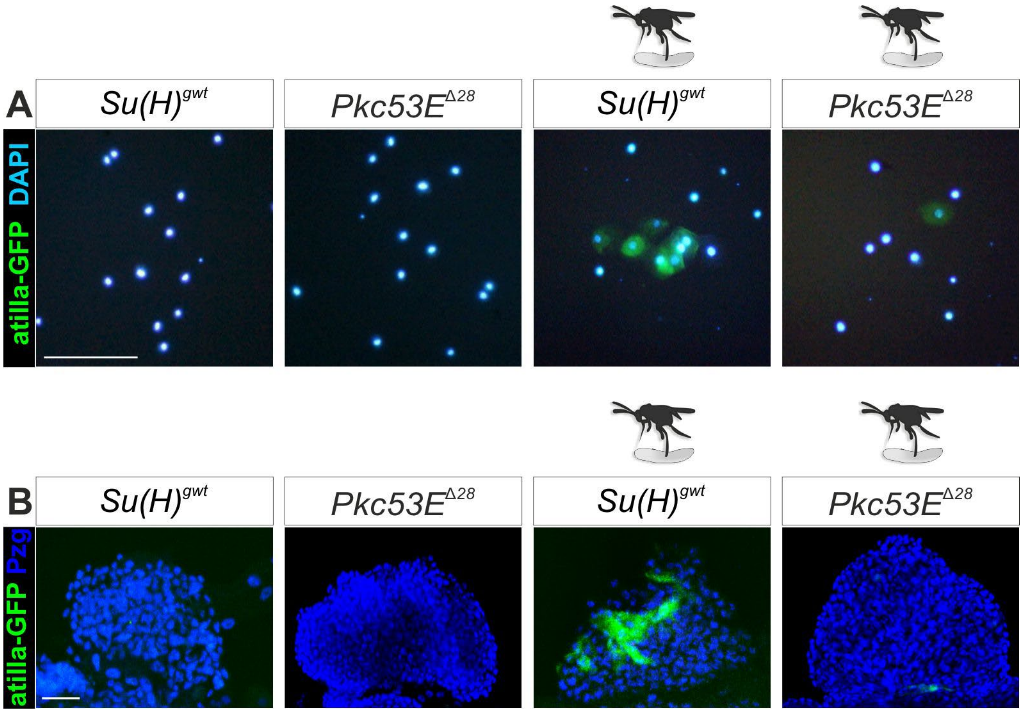

***Representative images for the various settings.***

**(A)** Representative images of hemolymph derived from larvae of the given genotype. Lamellocytes labelled green with *atilla*::GFP; nuclei blue (DAPI). **(B)** Representative images of one lymph gland 1° lobe of larvae with the given genotype. Lamellocytes are shown in green (*atilla*-GFP, nuclei in blue (anti-Pzg). (A,B) Infestation with *L. boulardi* indicated by the wasp schematic. Scale bar, 50 µm.

#### Tables Supplement

##### Figure 3 – supplement Table 1

List of kinases predicted to recognize S269 in Su(H) as substrate *in silico*

| Family of Kinase | Human | *Drosophila* |
|---|---|---|
| **AGC** | AKT1/AKT2 | Akt1 |
| | GRK5 | Gprk2 |
| | LATS1/LATS2 | Wts |
| | PDK1 | Pdk1 |
| | PRKCI | atypical PKC |
| | PRKCD | Pkc delta |
| | PRKG1 | For |
| | RPS61K3/RSKp90 | S6kII |
| **CAMK** | CAMK2A/CAMK2D | CamkII |
| | MAPK3 | Par-1 |
| | MAPKAPK3 | Mapk-Ak2 |
| | MYLK | Strn-Mlck |
| | PASK | Pask |
| | SIK2 | Sik2 |
| **CMGC** | CDK9/CycK/ CDK9/CycT | Cdk9 |
| | GSK3-alpha | Sgg |
| | GSK3-B | Gskt |
| **STE** | MAP2K3 | Lic |
| | MST1/MST2 | Hpp |
| | PAK4 | Mbt |
| **OPK** | AURA | AurA |
| | CHK1 | Grp |
| | DYRK1 | Mnb |
| | DYRK2 | Dyrk3 |
| | RAF1 | Raf |
| | WNK | Wnk |
| **Protein kinase like** | ATM | Tefu |
| | ATR | Mei-41 |
| | HASPIN | Haspin |
| | PRKB1 | Alc |

36 human candidate kinases, corresponding to 30 *Drosophila* kinases, were computationally predicted to have Su(H) S269 as a target phospho-site using GPS 3.0 software.

**Figure 3 – supplement Table 2**

List of kinases accepting the BTD domain of Su(H) as substrate *in vitro*

| Family of Kinase | Human | *Drosophila* |
|---|---|---|
| AGC | AKT1/AKT2/AKT3/SGK2 | Akt1 |
| | PKC alpha/PKC beta1/ PKC beta2/ PKC gamma | Pkc53E |
| | PKC delta | PKC delta |
| | PKC nu | Pkd |
| CAMK | CAMK1D/CAMK4 | CamkI |
| | CAMK2A/CAMK2B/ CAMK2D/CAMK2G | CamkII |
| | CHK2 | Lok |
| | MAPKAPK3 | MAPK-Ak2 |
| | SIK1/SIK2 | Sik2 |
| CkI | TTBK1 | Asator |
| CMGC | CDK1/CycA | Cdk1 |
| | CDK1/CycE CDK2/CycE CDK3/CycE CDK2/CycA | Cdk2 |
| | CDK7/CycH | Cdk7 |
| | CDK8/CycC | Cdk8 |
| | CLK2 | Doa |
| | GSK3-alpha | Sgg |
| | GSK3-B | Gskt |
| | JNK2 | Bsk |
| STE | CDC7/ASK | CG5790 |
| | MAP4K4 | Msn |
| | MAP4K5 | Hppy |
| | MEK1 | Dsor1 |
| | MST1/MST2 | Hpp |
| | NEK3/NEK9 | Niki |
| | STK39 | Fray |
| | TAOK2 | Tao |
| OPK | ACV-R2A | Put |
| | BMPR1A | Tkv |
| | B-RAF/B-RAFVE | Raf |
| | BUB1B | BubR1 |
| | CK2-alpha1/ CK2-alpha2 | CkII alpha |
| | HIPK1/HIPK2 | Hipk |
| | IRAK4 | Pll |
| | LIMK1/LIMK2 | Limk1 |
| | MAP3K1/MAP3K9 | Slpr |
| | PBK | CG8173 |
| | TLK2 | Tlk |
| | TSSK1 | CG14305 |
| | WEE1 | Wee1 |
| | WNK1/WNK2 | Wnk |

62 out of 245 human Ser/Thr kinases tested (equates to 25%) were identified to phosphorylate the BTD of Su(H) *in vitro*. These 62 human kinases correspond to 40 different kinases in *Drosophila*. 20 of the human kinases, corresponding to 10 *Drosophila* kinases were also predicted by the *in silico*

analysis (highlighted in green). The KinaseFinder Service was conducted by ProQinase (Freiburg Germany).

**Figure 3 – supplement Table 3**

Fly strains used for the larval crystal cell screen

| Kinase | Allele/RNAi/others | Source/Identity |
|---|---|---|
| Akt1 | $Akt^{04226}$/TM6ubi-GFP | Balancer changed from BL11627 (RRID:BDSC_11627) |
| Alc | $alc^{Ad2}$/CyO-GFP | Balancer changed from BL5510 (RRID:BDSC_5510) |
| Asator | TRIP_HMC04184attP2 dsRNA in Valium20 | BL55902 (RRID:BDSC_55902) |
| Bsk | TRIP_HMC03539attP2 dsRNA in Valium20 | BL53310 (RRID:BDSC_53310) |
| BubR1 | UAS-BubR1$^{DN}$ C-terminal truncated form | BL8380 (RRID:BDSC_8380) |
| | UAS-BubR1$^{DN}$ C-terminal truncated form | BL8382 (RRID:BDSC_8380) |
| CamkI | TRIP_JF02268attP2 dsRNA in Valium10 | BL26726 (RRID:BDSC_26726) |
| CamkII | TRIP_GL00237attP2/TM6B dsRNA in Valium22 | BL35330 (RRID:BDSC_26726) |
| | UAS-CamKII$^{T287A}$ Dominant negative version | BL29663 (RRID:BDSC_29663) |
| Cdk1 | $Cdk1^{E1-23}$/CyO-GFP | Balancer changed from BL6629 (RRID:BDSC_6629) |
| Cdk2 | TRIP_HM05163attP2 dsRNA in Valium10 | BL28952 (RRID:BDSC_28952) |
| Cdk7 | $Cdk7^{del}$/FM7-GFP | Balancer changed from BL4557 (RRID:BDSC_4557) |
| Cdk8 | TRIP_HMS05476attP40 dsRNA in Valium20 | BL67010 (RRID:BDSC_67010) |
| CG5790 | TRIP_HMJ23933attP40/CyO-GFP dsRNA in Valium20 | Balancer changed from BL62453 (RRID:BDSC_62453) |
| CG8173 | TRIP_JF01161attP2 dsRNA in Valium1 | BL31586 (RRID:BDSC_31586) |
| CG14305 | TRIP_HMC05158attP40 dsRNA in Valium20 | BL62151 (RRID:BDSC_62151) |
| CkII $\alpha$ | TRIP_JF01436attP2 dsRNA in Valium1 | BL31645 (RRID:BDSC_31645) |
| Doa | TRIP_HMC04193attP2 dsRNA in Valium20 | BL55908 (RRID:BDSC_55908) |
| | GD8588 dsRNA in pUASt | VDRC_19066 |
| Drak | $Drak^{del}$ | Neubuesser and Hipfner, 2010 |
| Dsor1 | TRIP_HMS00710 attP2/TM6B dsRNA in Valium20 | Balancer changed from BL32920 (RRID:BDSC_32920) |
| | KK102276 dsRNA in pUASt | VDRC_107276 |
| Dyrk3 | TRIP_HMC04155attP2 dsRNA in Valium20 | BL55882 (RRID:BDSC_55882) |
| Fray | TRIP_HMS01794attP2 dsRNA in Valium20 | BL38327 (RRID:BDSC_38327) |
| Gskt | TRIP_HMC05795attP2 dsRNA in Valium20 | BL64922 (RRID:BDSC_64922) |
| Hipk | TRIP_HMC05078attP40 dsRNA in Valium20 | BL60084 (RRID:BDSC_60084) |

| | | |
|---|---|---|
| Hpo | GD1570 dsRNA in pUASt | VDRC_7823 |
| | KK101704 dsRNA in pUASt | VDRC_104169 |
| Lic | GD7546 dsRNA in pUASt | VDRC_20166 |
| MAPk-Ak2 | TRIP_HMS04456attP40 dsRNA in Valium20 | BL57013 (RRID:BDSC_57013) |
| Mnb | KK102642 dsRNA in pUASt | VDRC_107066 |
| Msn | KK108948 dsRNA in pUASt | VDRC_101517 |
| | pValium20_attP40 shRNA | VDRC_330049 |
| Niki | TRIP_HMS01477attP2 dsRNA in Valium20 | BL35735 (RRID:BDSC_35735) |
| Par-1 | *par-1*$^{k06323}$/CyO-GFP | Balancer changed from BL10615 (RRID:BDSC_10615) |
| Pdk1 | KK108363 dsRNA in pUASt | VDRC_109812 |
| | TRIP_JF02807attP2 dsRNA in Valium10 | BL27725 (RRID:BDSC_27725) |
| Pkc53E | *Pkc53E*$^{\Delta28}$ | BL80988 (RRID:BDSC_80988) |
| | TRIP_JF02641 attP2 dsRNA in Valium10 | BL27491 (RRID:BDSC_27491) |
| | TRIP_HMS01195 attP2 dsRNA in Valium20 | BL34716 (RRID:BDSC_34716) |
| Pkc98E | TRIP_JF02470 attP2 dsRNA in Valium10 | BL29311 (RRID:BDSC_29311) |
| | TRIP_GL00174 attP2 dsRNA in Valium22 | BL35275 (RRID:BDSC_35275) |
| Pkd | *PKD*$^{cl4}$ | BL93864 (RRID:BDSC_93864) |
| | TRIP_JF03144attP2 dsRNA in Valium10 | BL28717 (RRID:BDSC_28717) |
| Pll | KK102624 dsRNA in pUASt | VDRC_103774 |
| Put | TRIP_JF02664 attP2 dsRNA in Valium10 | BL27514 (RRID:BDSC_27491) |
| Raf | *raf*$^{12}$/FM7-GFP | Balancer changed from BL5779 (RRID:BDSC_5779) |
| S6k | *S6k*$^{l-1}$/TM6B | BL32552 (RRID:BDSC_32552) |
| Sgg | *Sgg*$^{M1-1}$*/FM7-GFP* | Balancer changed from BL5402 (RRID:BDSC_5402) |
| Sik2 | TRIP_HMC04153 attP2 dsRNA in Valium20 | BL55880 (RRID:BDSC_55880) |
| Slpr | *slpr*$^{3P5}$*/ FM7-GFP* | Balancer changed from BL58795 (RRID:BDSC_58795) |
| | *slpr*$^{BS06}$*/ FM7-GFP* | Balancer changed from BL58807 (RRID:BDSC_58807) |
| | KK100726 dsRNA in pUASt | VDRC_106449 |
| | GD9771/CyO-GFP dsRNA in pUASt | VDRC_33518 |
| Tefu | TRIP_HMS02790 attP40 dsRNA in Valium20 | BL44073 (RRID:BDSC_44073) |
| Tkv | *tkv*$^{1}$ | BL427 (RRID:BDSC_427) |
| Wee | TRIP_HMC03331 attP40 dsRNA in Valium20 | BL51776 (RRID:BDSC_51776) |
| Wnk | TRIP_HMJ02087 attP40 dsRNA in Valium20 | BL42521 (RRID:BDSC_42521) |
| Gal4 driver/control lines | Allele/RNAi/other information | Source/Identity |
| *Hml*-Gal4 | Expresses Gal4 in lymph glands and circulating hemocytes | BL30141 (RRID:BDSC_30141) |
| *white*-RNAi | TRIP_JF01574attP2 dsRNA in Valium1 vector | BL31231 (RRID:BDSC_31231) |

**Figure 3 – supplement Table 4**

*Larval crystal cell screen*

Heated larvae of kinase mutants and/or *hml*-Gal4::UAS-kinase-RNAi/UAS-kinase$^{DN}$ genotypes were counted for the appearance of melanized crystal cells (cc) in the last two segments. If UAS-transgenes were used, the number of crystal cells was compared between uninduced (UAS-line alone) and induced with *hml*-Gal4. Mutant genotypes were related to the *Su(H)*$^{gwt}$ wild-type control.

Evaluation of cc numbers was ranged according to the following categories:

Dark blue: no cc present

Light blue: Less cc in comparison to controls and reference lines (at least 20%)

Beige: cc number within the range of *Su(H)*$^{gwt}$ and/or reference lines (up to ± 20% of reference)

Light green: cc number increase in the range of *Su(H)*$^{SA}$ (> 20-50% to reference)

Dark green: cc number increase above *Su(H)*$^{SA}$ level (> 50% to reference)

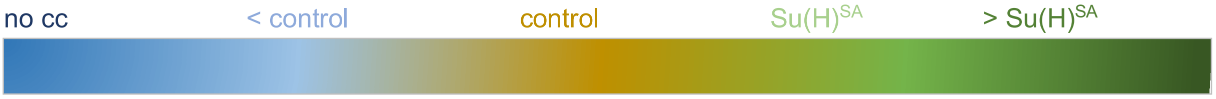

| Kinase (family) | Genotype | # Crystal cells | SD | Sample size (n) |
|---|---|---|---|---|
| Cdk1 (CMGC) | *cdk1*$^{E1-23}$ | 0.12 | 0.33 | 25 |
| Shaggy (CMGC) | *sgg*$^{M1-1}$ | 1.76 | 1.39 | 25 |
| Cdk7 (CMGC) | *cdk7*$^{del}$ | 71.68 | 17.52 | 44 |
| Raf (OPK) | *raf*$^{12}$ | 65.03 | 36.85 | 60 |
| Control | *Su(H)*$^{gwt}$ | 99.80 | 23.15 | 69 |
| Control | *hml-Gal4* | 110.32 (+11%) | 28.92 | 71 |
| Control | *white-RNAi* | 116.41 | 18.79 | 44 |
| | *hml::white-RNAi* | 110.67 (-5%) | 13.62 | 45 |
| Asator (CkI) | *Asator-RNAi* | 101.97 | 26.79 | 61 |
| | *hml::Asator-RNAi* | 106.86 (+5%) | 42.51 | 77 |
| CamKI (CAMK) | *CamkI*-RNAi | 84.96 | 35.42 | 51 |
| | *hml::CamkI*-RNAi | 76.44 (-11%) | 40.1 | 71 |
| Cdk2 (CMGC) | *cdk2-RNAi* | 133.26 | 29.39 | 54 |
| | *hml::cdk2-RNAi* | 123.11 (-8%) | 35.44 | 61 |
| CG5790 (STE) | *CG5790-RNAi* | 143.64 | 53.16 | 59 |
| | *hml::CG5790-RNAi* | 128.69 (-11%) | 32.96 | 67 |
| Dsor1 (STE) | *Dsor1-RNAi* (A) | 191.72 | 43.56 | 53 |
| | *hml::Dsor1-RNAi* | 159.04 (-17%) | 42.65 | 57 |
| | *Dsor1-RNAi* (B) | 151.86 | 34.92 | 56 |
| | *hml::Dsor1-RNAi* | 146.19 (- 4%) | 32.72 | 52 |
| Fray (STE) | *fray-RNAi* | 124.45 | 31.30 | 51 |
| | *hml::fray-RNAi* | 137.38 (+10%) | 40.13 | 65 |
| Hipk (OPK) | *hipk-RNAi* | 102.26 | 37.52 | 53 |
| | *hml::hipk-RNAi* | 121.35 (+19%) | 42.54 | 74 |
| Hpo (STE) | *hpo-RNAi* (A) | 166.13 | 44.05 | 48 |
| | *hml::hpo-RNAi* | 159.64 (- 4%) | 63.81 | 61 |
| | *hpo-RNAi* (B) | 74.91 | 25.85 | 55 |
| | *hml::hpo-RNAi* | 87.21 (+16%) | 42.58 | 62 |
| Lic (STE) | *lic-RNAi* | 80.53 | 29.37 | 45 |

| | | | | |
|---|---|---|---|---|
| | *hml::lic-RNAi* | 81.15 | 27.12 | 52 |
| MAPk-Ak2 (CAMK) | *MAPk-Ak2-RNAi* | 104.09 | 25.96 | 56 |
| | *hml::MAPk-Ak2-RNAi* | 117.98 (+13%) | 31.27 | 52 |
| Mnb (OPK) | *mnb-RNAi* | 116.10 | 42.25 | 40 |
| | *hml::mnb-RNAi* | 115.40 | 30.82 | 43 |
| Sik2 (CAMK) | *Sik2-RNAi* | 142.38 | 28.81 | 47 |
| | *hml::Sik2-RNAi* | 151.47 (+6%) | 34.80 | 53 |
| Tefu (Protein kinase like) | *tefu-RNAi* | 56.69 | 18.12 | 55 |
| | *hml::tefu*-RNAi | 56.41 | 28.15 | 61 |
| Wee1 (OPK) | *Wee1-RNAi* | 105.44 | 29.4 | 55 |
| | *hml::Wee1-RNAi* | 121.37 (+15%) | 36.43 | 68 |
| Reference | *Su(H)*$^{SA}$ | 135.77 (+36%) | 25.99 | 70 |
| Akt (AGC) | *akt1*$^{04226}$ | 135.1 (+36%) | 43.14 | 48 |
| BubR1 (OPK) | *BubR1*$^{DN}$ (A) | 77.92 | 20.09 | 52 |
| | *hml::BubR1*$^{DN}$ | 105.14 (+35%) | 22.25 | 58 |
| | *BubR1*$^{DN}$ (B) | 84.78 | 25.43 | 49 |
| | *hml::BubR1*$^{DN}$ | 112.85 (+33%) | 34.11 | 48 |
| Bsk (CMGC) | *bsk-RNAi* | 110.23 | 41.19 | 62 |
| | *hml::bsk-RNAi* | 149.03 (+35%) | 35.84 | 71 |
| CamkII (CAMK) | *CamkII*$^{DN}$ | 163.97 | 30.53 | 58 |
| | *hml:: CamkII*$^{DN}$ | 221.35 (+35%) | 54.03 | 46 |
| | *CamkII-RNAi* | 152.96 | 49.05 | 50 |
| | *hml:: CamkII-RNAi* | 188.85 (+23%) | 61.87 | 68 |
| Cdk8 (CMGC) | *cdk8-RNAi* | 115.63 | 29.36 | 57 |
| | *hml::cdk8-RNAi* | 170.63 (+47%) | 31.85 | 56 |
| CG14305 (OPK) | *CG14305-RNAi* | 96.49 | 27.27 | 43 |
| | *hml::CG14305-RNAi* | 134.55 (+39%) | 36.37 | 65 |
| Doa (CMGC) | *Doa-RNAi (A)* | 143.35 | 22.56 | 63 |
| | *hml::Doa-RNAi* | 174.02 (+21%) | 36.58 | 60 |
| | *Doa-RNAi (B)* | 114.20 | 23.34 | 46 |
| | *hml::Doa-RNAi* | 170.42 (+49%) | 35.73 | 45 |
| Dyrk3 (OPK) | *Dyrk3-RNAi* | 105.07 | 27.65 | 60 |
| | *hml::Dyrk3-RNAi* | 151.95 (+45%) | 35.43 | 59 |
| Gskt (CMGC) | *gskt-RNAi* | 98.60 | 35.01 | 50 |
| | *hml::gskt-RNAi* | 137.46 (+39%) | 33.41 | 63 |
| Msn (STE) | *msn-RNAi* (A) | 82.83 | 36.11 | 53 |
| | *hml::msn-RNAi* | 122.58 (+48%) | 47.36 | 53 |
| | *msn-RNAi* (B) | 102.79 | 26.29 | 63 |
| | *hml::msn-RNAi* | 135.92 (+32%) | 34.99 | 62 |
| Niki (STE) | *niki-RNAi* | 120.87 | 29.91 | 47 |
| | *hml::niki-RNAi* | 147.0 (+22%) | 53.59 | 51 |
| Pdk1 (AGC) | *Pdk1-RNAi* (A) | 130.33 | 30.34 | 48 |
| | *hml::Pdk1-RNAi* | 188.52 (+45%) | 33.82 | 50 |
| | *Pdk1-RNAi* (B) | 123.83 | 30.06 | 46 |
| | *hml:: Pdk1-RNAi* | 168.24 (+47%) | 28.89 | 46 |
| Pkc53E (AGC) | *Pkc53E*$^{\Delta 28}$ | 137.40 (+38%) | 47.92 | 70 |
| | *PKc53E-RNAi* (A) | 140.42 | 25,75 | 55 |
| | *hml::Pkc53E-RNAi* | 185.09 (+32%) | 34.26 | 70 |
| | *PKc53E-RNAi* (B) | 111.21 | 33.79 | 52 |
| | *hml::Pkc53E-RNAi* | 163.29 (+47%) | 34.59 | 70 |
| Pkc98E (AGC) | *Pkc98E-RNAi* (A) | 81.03 | 35.02 | 60 |
| | *hml::Pkc98E-RNAi* (A) | 111.52 (+38%) | 40.73 | 54 |
| | *Pkc98E-RNAi* (B) | 126.25 | 30.84 | 61 |
| | *hml::Pkc98E-RNAi* (B) | 165.67(+31%) | 29.49 | 85 |
| PKD (AGC) | *PKD*$^{cl4}$ | 137.23 (+38%) | 31.59 | 60 |
| | *PKD-RNAi* | 125.85 | 30.45 | 47 |
| | *hml::PKD-RNAi* | 161.18 (+28%) | 48.40 | 60 |
| Pll (TKL) | *pll-RNAi* | 125.76 | 41.70 | 55 |

| | | | | |
|---|---|---|---|---|
| | *hml::pll-RNAi* | 162.76 (+29%) | 42.52 | 55 |
| Put (Protein kinase like) | *put-RNAi* | 129.04 | 28.03 | 50 |
| | *hml::put-RNAi* | 176.02 (+36%) | 41.26 | 45 |
| S6K (AGC) | *S6k*$^{l-1}$/+ (no hz) | 145.1 (+45%) | 36.07 | 58 |
| Wnk (OPK) | *Wnk-RNAi* | 89.89 | 26.25 | 70 |
| | *hml::Wnk-RNAi* | 127.10 (+41%) | 29.81 | 70 |
| Alc (Protein kinase like) | *alc*$^{Ad2}$/+ (no hz) | 158.75 (+59%) | 54.65 | 52 |
| CG8173 (OPK) | *CG8173-RNAi* | 57.98 | 17.70 | 58 |
| | *hml::CG8173-RNAi* | 114.85 (+98%) | 26.13 | 53 |
| CkIIα (OPK) | *CkIIα-RNAi* | 86.43 | 27.18 | 63 |
| | *hml::CkIIα-RNAi* | 169.7 (+96%) | 34.86 | 63 |
| Drak (CAMK) | *drak*$^{del}$ | 151.62 (+52%) | 39.43 | 61 |
| Par-1 (CAMK) | *par-1*$^{K06323}$/+ (no hz) | 151.6 (+52%) | 48.49 | 58 |
| Slpr (STE) | *slpr*$^{B506}$/*slpr*$^{3P5}$ | 155.08 (+56%) | 36.12 | 50 |
| | *slpr-RNAi* (A) | 49.64 | 16.91 | 47 |
| | *hml:: slpr-RNAi* | 101.32 (+104%) | 29.55 | 50 |
| | *slpr-RNAi* (B) | 106.69 | 25.09 | 59 |
| | *hml:: slpr-RNAi* | 170.76 (+60%) | 26.57 | 59 |
| Tkv (OPK) | *tkv*$^{1}$ | 173.46 (+74%) | 43.04 | 54 |

#### SUPPLEMENTAL TABLE S1 KEY RESOURCES

| REAGENT or RESOURCE | SOURCE | IDENTIFIER |
|---|---|---|
| Antibodies and sera | | |
| mouse monoclonal anti-Hnt, 1G9 developed by H. Lipshitz | Developmental Studies Hybridoma Bank | RRID: AB_528278 |
| mouse monoclonal anti-mys, CF.6G11 developed by D. Brower | Developmental Studies Hybridoma Bank | RRID: AB_528310 |
| mouse monoclonal anti-beta tubulin, E7 developed by M. Klymkowsky | Developmental Studies Hybridoma Bank | RRID: AB_2315513 |
| mouse monoclonal anti-myc 9B11 | Cell Signaling Techn. | RRID: AB_331783; Cat#2276 |
| mouse monoclonal anti-Hemese | Kurucz et al., 2003; gift from I. Andó, Szeged, Hungary | N/A |
| guinea pig polyclonal anti-Pzg | Kugler and Nagel, 2007 | N/A |
| mouse monoclonal anti-PPO1, 12F6 | Trenczek, Bennich, 1992; gift from T.E.Trenczek, Giessen, Germany | N/A |
| mouse monoclonal anti-GST 8-326rabbit polyclonal anti-pS269 | this work, DAVIDS Biotechnology GmbH | N/A |
| rabbit polyclonal anti-GFP | Santa Cruz | RRID: AB_641123; Cat#sc-8334 |
| rabbit polyclonal anti-mCherry | GeneTex | RRID: AB_2721247; Cat#GTX128508 |
| rat monoclonal anti-HA 3F10 | ROCHE | RRID:AB_390918 Cat#11867423001 |
| donkey polyclonal anti-mouse IgG, **Cy$^{TM}$3** | Jackson Immuno-Research Laboratories | RRID: AB_2315777 Cat#715-165-151 |
| donkey polyclonal anti-mouse IgG**, Cy$^{TM}$5** | Jackson Immuno-Research Laboratories | RRID: AB_2340820 Cat#715-175-151 |
| donkey polyclonal anti- guinea pig IgG, **Cy$^{TM}$5** | Jackson Immuno-Research Laboratories | RRID: AB_2340462 Cat#706-175-148 |
| donkey polyclonal anti- rabbit IgG, **FITC** | Jackson Immuno-Research Laboratories | RRID: AB_2315776 Cat#711-095-152 |
| goat polyclonal anti-guinea pig IgG, **Alexa Fluor® 647** | Jackson Immuno-Research Laboratories | RRID: AB_2337446; Cat#106-605-003 |
| goat polyclonal anti-mouse IgG, **FITC** | Jackson Immuno-Research Laboratories | RRID: AB_2338601; Cat#1115-095-166 |
| goat polyclonal anti-rabbit IgG, alkaline phosphatase | Jackson Immuno-Research Laboratories | RRID: AB_2337947; Cat#111-055-003 |
| goat polyclonal anti-rat IgG, alkaline phosphatase | Jackson Immuno-Research Laboratories | RRID: AB_2338148; Cat#112-055-003 |
| goat polyclonal anti-mouse IgG, alkaline phosphatase | Jackson Immuno-Research Laboratories | RRID: AB_2338528; Cat#115-055-003 |
| normal donkey serum | Jackson Immuno-Research Laboratories | RRID: AB_2337258 Cat#017-000-121 |
| normal goat serum | Jackson Immuno-Research Laboratories | RRID: AB_2336990 Cat#005-000-121 |
| Chemicals, peptides, and recombinant proteins | | |
| DAPI | Cell Signaling Techn. | Cat#4083 |
| DNase I | New England Biolabs GmbH | Cat#M0303 |
| PMA, Phorbol-12-myristat-13-acetat | Sigma-Aldrich | Cat#P8139-1MG |

| | | |
|---|---|---|
| Staurosporine | Sigma-Aldrich | Cat#S4400-1MG |
| Protease inhibitor cocktail, cOmplete$^{TM}$ ULTRA-tablets Mini | Roche | Cat# 5892791001 |
| Wheat germ agglutinin (WGA), Alexa Fluor$^{TM}$ 647 conjugate | Fisher Scientific | Cat# 11510826 |
| Phalloidin, coupled to rhodamine | Invitrogen, Thermo Fisher | Cat# R415 |
| Vectashield mounting medium | Biozol | Cat#VEC-H_1000 |
| RFP-Trap Magnetic Agarose | ChromoTek | Cat#rtma-20 |
| Amylose resin | New England Biolabs GmbH | Cat#E8021S |
| activated PKC$\alpha$ | ProQinase | Cat#0222-0000-1 |
| activated Akt1 | ProQinase | Cat#1379-0000-2 |
| activated GSK3 beta | ProQinase | Cat#0310-0000-1 |
| activated S6K | ProQinase | Cat#0318-0000-2 |
| CAMK2D | Invitrogen | NP_742113 |
| Pseudosubstrate RFARLGSLRQKNV | peptides & elephants | PS |
| Su(H) peptide ALFNRLRSQTVSTRY | peptides & elephants | $S^{wt}$ |
| Su(H)SA peptide ALFNRLRAQTVSTRY | peptides & elephants | $S^{SA}$ |
| Su(H) phosphopeptide NLRLpSQTVSTRYLHVE | DAVIDS Biotechnology GmbH | N/A |
| PhosSTOP™ (Phosphatase inhibitor) | Roche | Cat#4906837001 |
| Critical commercial assays and tools | | |
| ADP-Glo$^{TM}$ Kinase Assay | Promega | Cat#V6930 |
| PolyATract System Kit 1000 | Promega | Cat#Z5400 |
| Dynabeads$^{TM}$ mRNA DIRECT micro purification kit | Invitrogen, Thermo Fisher | Cat#61021 |
| qScriber$^{TM}$ cDNA Synthesis Kit | highQu | Cat#RTK0104 |
| Q5® Site directed Mutagenesis Kit | New England Biolabs GmbH | Cat# E0554S |
| Blue S'Green qPCR Kit | Biozym | Cat#331416 |
| Dual-Luciferase® Reporter Assay | Promega | Cat# E1910 |
| PAP PEN | Kisker Biotech | Cat# MKP-1 |
| Experimental models: Cell lines | | |
| *RBPj*$^{KO}$ HeLa cells | Wolf *et al.,* 2019 | N/A |
| Experimental models: Organisms/strains | | |
| Parasitoid wasps | | |
| *Leptopilina boulardi* | B. Häußling, J. Stökl, Bayreuth | N/A |
| *Leptopilina heterotoma* | B. Häußling, J. Stökl, Bayreuth | N/A |
| *Asobara japonica* | B. Häußling, J. Stökl, Bayreuth | N/A |
| *Drosophila melanogaster* stocks | | |
| *atilla*-GFP, i.e. $w^{1118}$; Mi{ET1}$atilla^{MB03539}$ | Bloomington Drosophila Stock Center | BDSC:23540 |
| *He*-Gal4 UAS-GFP, i.e. $w^{*}$; P{He-GAL4.Z}85, P{UAS-GFP.nls}8 | Bloomington Drosophila Stock Center | BDSC:8700 |
| *hml*-Gal4, i.e. $w^{1118}$; P{Hml-GAL4.Δ}3 /MKRS | Bloomington Drosophila Stock Center | BDSC:30141 |
| *hmlΔ*-Gal4 UAS-GFP, i.e. $w^{1118}$; P{Hml-GAL4.Δ}3, P{UAS-2xEGFP}AH3/MKRS | Bloomington Drosophila Stock Center | BDSC:30142 |

| | | |
|---|---|---|
| *PPO3*-Gal4 UAS mCD8-GFP | Dudzic *et al.*, 2015 | N/A |
| *lz*-Gal4 | Lebestky *et al.*, 2000 | N/A |
| *NRE-GFP, i.e.* $w^{1118}$; P{NRE-EGFP.S}1 | Bloomington Drosophila Stock Center | BDSC:30728 |
| *Pkc53E-EGFP, i.e. Pkc53*$E^{MI05296\text{-}GFSTF.0}$ | Bloomington Drosophila Stock Center | BDSC:59413 |
| UAS-*white-RNAi, i.e.* $y^1$ $v^1$; P{TRiP.JF01574}attP2/TM3, $Ser^1$ | Bloomington Drosophila Stock Center | BDSC:31231 |
| UAS-*N-RNAi, i.e.* P{UAS-N.RNAi.P}14E, $w^*$ | Bloomington Drosophila Stock Center | BDSC:7078 |
| UAS-*sgRNA-Pkc53E* | Vienna Drosophila Resource Center | VDRC341127 |
| UAS-*µMCas9* | Vienna Drosophila Resource Center | VDRC 340002 |
| UAS-HA-*Pkc53E* | this study | N/A |
| *vasa-φC31*; *96E-attB*/TM3 | Bischof *et al.*, 2007 | N/A |
| *Su(H)*$^{gwt}$ | Praxenthaler *et al.*, 2017 | N/A |
| *Su(H)*$^{gwt\text{-}mCh}$, *i.e.* $y^1$ $w^*$; TI{TI}Su(H)$^{gwt\text{-}mCh}$ | Praxenthaler *et al.*, 2017 | BDSC:94607 |
| *Su(H)*$^{S269A}$, *i.e.* $y^1$ $w^*$; TI{TI}Su(H)$^{S269A}$ | Frankenreiter *et al.*, 2021 | BDSC:94609 |
| *Su(H)*$^{S269D}$/*CyO-GFP, i.e.* $y^1$ $w^*$; TI{TI}Su(H)$^{S269D}$/CyO, P{GAL4-Hsp70.PB}TR1, P{UAS-GFP.Y}TR1 | Frankenreiter *et al.*, 2021 | BDSC:94610 |
| *Su(H)*$^{S269A\text{-}mCh}$ | this study | N/A |
| Kinase mutant flies tested in the larval kinase screen are listed in Figure 3 - supplemental Table 3 | | |
| Oligonucleotides | | |
| See supplemental Table S2, Oligonucleotides | | |
| Recombinant DNA | | |
| *Pkc53E* cDNA | Drosophila Genome Resource Center | GH03188 |
| *Pkc53E*$^{EDDD}$ | this work | N/A |
| Su(H) cDNA | Maier *et al.*, 2011 | N/A |
| Su(H) BTD | this work | N/A |
| 2xMyc-Su(H) | this work | N/A |
| HSV-TK 2xMyc-Su(H)-VP16 | this work | N/A |
| pGL3 NRE | Bray *et al.*, 2005 | N/A |
| pUAST Su(H)-VP16 | Cooper *et al.*, 2000 | N/A |
| pUAST-attB | Bischof *et al.*, 2007 | RRID:DGRC_1419 |
| pBT-3xHA | this work | N/A |
| pGEX-2T | Smith and Johnson, 1988 | N/A |
| pMAL | Riggs, 1994 | N/A |
| pCDNA3.1 | Invitrogen | Cat#V79020 |
| pRL TK | Promega | Cat#E2241 |
| Software and algorithms | | |
| BoxPlotR | Spitzer *et al.*, 2014 | http://shiny.chemgrid.org/boxplotr/ |
| GPS3.0 software | Xue *et al.*, 2011 | |
| *ImageJ* 1.51 | Schindelin *et al.*, 2012 | https://imagej.nih.gov/ij/ |

| | | |
|---|---|---|
| GraphPad Prism version 9.0 | Graphpad Software, Inc. | https://www.graphpad.com/ |
| MIC PCR software version v2.12.7 | bms/Biozym | Cat#68MiC-HRM |
| Weka machine learning and data analysis software version 3.8 | Eibe *et al.*, 2016 | https://waikato.github.io/weka-site/index.html |

*Newly generated materials are available upon request.*

**SUPPLEMENTAL TABLE S2 Oligonucleotides**

| Primer | Sequence 5‘ -> 3‘ | Purpose | PCR conditions |
|---|---|---|---|
| ***Su(H) cloning and verification*** | | | |
| Su(H) BTD_UP | AAG GGA TCC TCG CTA AAG AAT GCC GAT CTG TG | BTD-amplification with 5’ *Bam*HI and 3’ *Eco*RI sites for subcloning into pGEX-2T | 52.5°C Annealing; Product length: 525 bp |
| Su(H) BTD_LP | CAT GAA TTC TCA GAA CTG GTA CTC AGC CTT GTC GG | | |
| Myc-Tag_UP | AAT TCA TGG AGC AGA AGC TGA TCT CGG AGG AGG ATC TAG AGC AGA AGC TGA TCT CGG AGG AGG ATC TAG AGC | | Addition of myc-tag to Su(H) cDNA by insertion of annealed primers via *Eco*RI |
| Myc-Tag_LP | AAT TGC TCT AGA TCC TCC TCC GAG ATC AGC TTC TGC TCT AGA TCC TCC TCC GAG ATC AGC TTC TGC TCC ATG | | |
| attR UP | CGG GGG ATC CAC TAG TTC TAG ATG TAG G | Verification of *Su(H)* genome engineered flies | 57°C Annealing; Product length: 851 bp |
| Su(H) attR LP | CGC GCA TAG TTG TGC TCC CTG TTC G | | |
| 5‘Exon4 | TTA TGC CAC CGG TCT GAC CTT CA | Verification of *Su(H)*-mCherry tag | 60°C Annealing; Product length: 1344 bp |
| Su(H) flox LP | CAA CCG CAT CTA AAA ATC GTC TAT AAA CTT ACA TC | | |
| 5’Exon2 | CCC AAA AGT CCT ATG GCA ATG A | Differentiation of $Su(H)^{gwt}$ versus $Su(H)^{S269A}$ via *Alw44*I digest | 59°C Annealing; Product length: 1.5kb in wild type vs 1.1+0.41 in mutant |
| Su(H) hom-reg LP | GGA TAA GCC GCT ACC ATG ACT ATT | | |
| ***Pkc53E* cloning, verification and mutagenesis** | | | |
| 3HA-Tag_UP *Acc65*I/*Xho*I | GTA CCA TGT ATC CCT ATG ATG TGC CAG ACT ATG CTG GCT ATC CAT ATG ATG TTC CTG ATT ATG CTG GAT ACC CTT ATG ATG TGC CAG ACT ATG CCC | | Generation of pBT -3HA by insertion of annealed primers into Acc65I/XhoI of pBT |
| 3HA-Tag_LP *Acc65*I/*Xho*I | TCG AGG GCA TAG TCT GGC ACA TCA TAA GGG TAT CCA GCA TAA TCA GGA ACA TCA TAT GGA TAG CCA GCA TAG TCT GGC ACA TCA TAG GGA TA**C** ATG | | |
| *Xho*I_Pkc53E_UP | TTT ACT CGA GAT GTC GGA GGG CAG | Amplification of Pkc53E cDNA with 5’ *Xho*I and 3’ *Xba*I sites for subcloning into pBT-3HA vector | 59°C Annealing; Product length: 2.1 kb |
| *Xba*I_Pkc53E_LP | CTG ATC TAG ACT ATG GGC TGA AAA CAT ATT CG | | |
| *Pkc53E Exon1_UP* | CGG AGG GCA GCG ATA ACA ACG G | Identification of Pkc53E null mutant | 59°C Annealing; Product length: |

| | | | |
|---|---|---|---|
| *Pkc53E Exon1_LP* | CAA GTG CCG TGG AGG GAA GTG GG | | 553 kb in wild type, none in mutant |
| *Pkc53E_A34E_UP* | CCG CAA AGG **AGA G**CT CAA GAA GAA G | Site directed mutagenesis *A34E* | 61°C Annealing |
| *Pkc53E_A34E_LP* | AGG CGG GAC TTC ATT TTG | | |
| *Pkc53E_T508D_UP* | CGG TAC CCC TGA TTA CAT TGC TCC AG | Site directed mutagenesis *T508E* | 60°C Annealing |
| *Pkc53E_ T508D_LP* | CAG AA**A TC**C TTT GTG GTC TTA TCA CC | | |
| *Pkc53E_T650E_UP* | **GAT** CCC ACG GAC AAG GTG TTT ATG | Site directed mutagenesis *T650E* | 58°C Annealing |
| *Pkc53E_ T650E_LP* | CAG ATC TGT TTT CTC TGA TGT GAA CTG | | |
| *Pkc53E_S669D_UP* | AAT CCC GAG TAT GTT TTC AGC CCA TAG TC | Site directed mutagenesis *S669D* | 61°C Annealing |
| *Pkc53E_ S669D_LP* | CAT GTA **GTC** GAA GCC AAC GAA TTC CGA C | | |
| ***Pkc53E RT-PCR and qRT-PCR*** | | | |
| Pkc53E_RT-PCR UP | CTG CGG TAC ACC TGA TTA CAT TGC TC | RT-PCR for Pkc53E transcript | 58°C Annealing; Product length: 430 bp |
| Pkc53E_RT-PCR LP | CGT GGG CGT CAA GTC TGT TTT CT | | |
| Tub56D_229 UP | GAA CCT ACC ACG GTG ACA GCG A | RT-PCR for *Tubulin56D* transcript | 65°C Annealing; Product length: 299 bp |
| Tub56D_507 LP | GAA GCC AAG CAG GCA GTC GCA | | |
| GFP_1 UP | GCC ACA AGT TCA GCG TGT CCG | qRT-PCR for NRE-GFP transcript | |
| GFP_1 Rev | GTA GGT CAG GGT GGT CAC GAG GG | | |
| atilla_4 UP | AAA CAA GTG ATT TTC GTG CTC CT | qRT-PCR for *atilla* transcript | PP7306 |
| atilla_4 Rev | CGC GGA TGT TAG AGG CAG A | | |
| cyp33_2 UP | CTC TGC GGA CGC ACA ATT C | qRT-PCR for *cyp33* transcript | PP14577 |
| cyp33_2 Rev | TGC AAC CAG TCG TCA TCT GC | | |
| Tbp_1 UP | TAA GCC CCA ACT TCT CGA TTC C | qRT-PCR for *Tbp* transcript | PP1556 |
| Tbp_1 Rev | GCC AAA GAG ACC TGA TCC CC | | |
| ***genotyping/verification*** | | | |
| Gal4 UP | CCG TCA CAG ATA GAT TGG CTT | Presence of Gal4 driver | 55°C Annealing; Product length: 677 bp |
| Gal4 LP | AAG ATG TAG GGC TGT CAC CAA | | |
